# Protocol for spatial characterization of ECM collagen-GAG in CRC tumor microenvironment

**DOI:** 10.1101/2025.06.25.661355

**Authors:** Libby A. Boykin, Sneha Jayashankar, Khidr Kishan K. Budhwani, Brahma Mubarak K. Budhwani, Diya Samal, Chelsea L. Crawford, Allan Tsung, Karim I. Budhwani

**Affiliations:** CerFlux, Birmingham, AL, USA; Department of Surgery, University of Virginia School of Medicine, Charlottesville, VA, USA

**Keywords:** Alcian blue, picrosirius red, dual-stain protocol, extracellular matrix (ECM), tumor microenvironment (TME), collagen, glycosaminoglycans (GAGs), colorectal cancer (CRC)

## Abstract

The extracellular matrix (ECM) plays a critical role in colorectal cancer (CRC) progression and therapeutic resistance. Accurate characterization of ECM composition and architecture is essential for understanding how CRC evades therapy, yet most protocols either assess ECM components in isolation or remain technically challenging. Here we present a robust yet simple protocol for spatial characterization of collagen and glycosaminoglycan (GAG) organization within the CRC tumor microenvironment (TME). Our method combines Alcian Blue and Picrosirius Red staining procedures with standardized tissue processing and imaging protocols. The protocol enables simultaneous visualization, assessment, and quantification of collagen and GAG distribution patterns in formalin-fixed, paraffin-embedded tissue sections. Key methodological advances include optimized dual-staining approach with distinct blue-red coloration for straightforward spectral separation on digital imaging systems, standardized reagent preparations, and validated imaging parameters. The complementary wavelengths facilitate both visual interpretation and potential digital separation, offering advantages over multi-component stains with overlapping spectral ranges. Validation across multiple CRC patient specimens demonstrates excellent reproducibility with consistent staining intensity using standard histology equipment. Because the resulting spatial maps can be compared directly with engineered or *ex vivo* tumor models, the protocol also provides a practical benchmark for microenvironment validation. This standardized approach to ECM visualization will advance TME research, support morphological studies, and enable comparative analyses across CRC subtypes. The methodology can be adapted to other solid tumor types and integrated with complementary techniques including digital pathology workflows for comprehensive microenvironment characterization and enhanced analysis capabilities.

## 1. INTRODUCTION

Colorectal cancer (CRC) continues to represent a substantial public health burden worldwide, ranking 3^rd^ in incidence and 2^nd^ in cancer mortality with an estimated 1.9 million new cases and 935,000 deaths annually.^1^ Of particular concern is the rising incidence of CRC in younger adult populations. In the USA and other high-income countries, rates among individuals under the age of 50 have increased by ∼2% per year in recent years.^2^ Consequently, the disease burden is shifting demographically, with approximately 20% of CRC diagnoses now occurring in patients younger than 55.^2^ Because younger patients are often diagnosed before they reach the current recommended age for screening, early-onset cases frequently present at advanced stages. CRC has now become the leading cause of cancer-related death in men under 50 (and second in women), marking a significant departure from mortality patterns observed in the late 1990s.^3^ These patterns underscore a worsening outlook for young-onset CRC, reversing earlier progress in overall survival rates and emphasizing the critical need for improved understanding and management of the disease.^2^

Emerging research implicates the extracellular matrix (ECM) of the tumor microenvironment (TME) as a key driver of CRC progression, metastasis, and clinical outcomes. The ECM is an intricate network of macromolecules that give the tissue structure and organization. Components of the ECM can link together to give complex mechanical properties to the tissue. CRC tumors typically trigger a desmoplastic stromal reaction enriched in fibrillar collagens and glycosaminoglycans (GAGs) such as hyaluronic acid (HA), which in turn results in an abnormally stiff TME.^4^ This remodeled ECM promotes invasion further as cancer-associated fibroblasts produce abundant collagen and proteoglycans, increase matrix rigidity, and create tracks that facilitate cancer cell migration and dissemination.^4^ Collagen is the predominant structural protein in the TME that profoundly influences tumor behavior, and aberrant collagen remodeling in CRC has been linked to greater invasiveness and poorer survival.^5^ GAGs are mucopolysaccharides or negatively charged groups of polysaccharides formed from a chain of disaccharides. GAGs have been implicated in both increasing and decrease cancer cell proliferation.^6^ An HA-rich stroma promotes an immunosuppressive, pro-tumor microenvironment. High levels of HA in the TME and overexpression of its receptor RHAMM correlate with advanced disease and adverse prognosis in CRC.^7^ Collectively, the ECM modulates disease progression and also serves as a key prognostic indicator.

TME ECM is also implicated in treatment resistance. Dense collagen networks can impede drug delivery and foster protective niches for cancer cells, while biomechanical and biochemical cues from the ECM activate survival pathways that blunt therapeutic efficacy.^1^ These protective effects potentially contribute to persistent translational gaps between preclinical models and clinical outcomes. Such gaps widen the “*valley of death*” for new cancer drugs, where over 90% of drugs that prove effective in animal studies fail to translate in humans, often due to species differences in TME and drug response.^8,9^ Accordingly, both the US Food and Drug Administration (FDA) and the National Institutes of Health (NIH) have called for a paradigm shift in translational cancer research toward human-relevant with *new approach methodologies* (NAMs)^10^ and *Novel Alternative Methods* (NAMs)^11^ respectively. Regulatory agencies have begun phasing out traditional animal models in drug development, encouraging the adoption of human-derived 3D models (patient organoids, *ex vivo* tissue cultures, organ-on-a-chip systems) and other NAMs that better recapitulate human tumor biology.^12^ These innovative platforms have shown promise in predicting patient drug responses,^13^ but their success hinges on authentically recreating features like the tumor ECM. There is thus a pressing need for methodologies that characterize and incorporate the human CRC ECM to drive more predictive preclinical models.^12^

In this context, accurate spatial characterization of the CRC ECM is paramount. Clinically, mapping the distribution and organization of collagen and GAGs in tumors could yield novel stromal biomarkers to refine prognosis and track disease progression at the bedside.^1,5^ Concurrently, translating such detailed ECM insights to the laboratory will guide the development and validation of NAMs, ensuring these next-generation models faithfully mimic the matrix conditions that influence tumor behavior and drug response.^12^ Here we present a dual-staining protocol using Alcian Blue and Picrosirius Red on formalin-fixed, paraffin-embedded (FFPE) CRC tissue sections to spatially delineate collagen and GAG/HA content and architecture in the TME. By capturing the nuanced ECM landscape of CRC TME, this approach aims to support prognostic assessments in patients and accelerate creation of human-relevant preclinical models for CRC drug discovery.

Alcian Blue is a cationic dye that stains and identifies negatively charged acidic mucosubstances, particularly acidic polysaccharides such as GAGs. Chemically, it is a copper-containing phthalocyanine dye (C56H70Cl4CuN16S4, MW ∼1300 g/mol) with a planar structure bearing four covalently attached isothiouronium groups, each carrying a positive charge.^14^ This highly basic structure enables Alcian Blue to form strong electrostatic (ionic) bonds with negatively charged molecules in tissue. Its four cationic centers bind specifically to anionic functional groups on acidic mucopolysaccharides, anchoring the dye to its targets. These ionic “salt linkages” are what keep the stain bound to tissue components during subsequent washing and staining steps, preventing the dye from being washed off easily. Staining intensity is proportional to the density of negative charges, as areas with high concentrations of acidic groups bind more dye and appear more intensely blue. Because these interactions are electrostatic, staining specificity can be modulated by pH. At mildly acidic pH (∼2.5), it stains both sulfated and carboxylated acidic mucopolysaccharides blue, while at lower pH (∼1.0), only sulfated GAGs bind the dye since carboxyl groups become protonated and lose their charge. This pH-dependent binding allows differentiation of subtypes of acidic glycoconjugates based on their charge densities.^15^

The other stain of interest is Picrosirius Red which consists of Sirius Red F3B (Direct Red 80), a poly-azo dye, dissolved in picric acid (hence the name picrosirius). This large planar molecule (C45H26N10Na6O21S6, MW ≈1373 g/mol)^16^ contains four azo bonds and six sulfonate groups that provide a strong natural affinity to collagen,^17^ allowing it to bind very easily to collagen fibers even when dissolved in picric acid.^18^ This is particularly important in CRC tumors due to the abundance of collagen in the TME. The dye binds specifically to basic amino acid residues such as lysine, hydroxylysine, and arginine in collagen through multipoint electrostatic interactions.^17^ Picric acid creates an acidic environment (∼pH 2) that protonates these amino groups, enhancing the binding affinity between the dye and the collagen in the TME. Picrosirius Red molecules attach along the length of collagen fibrils via multipoint interactions wherein the sulfonate groups engage in ionic bonds while the extended planar structure allows for hydrogen bonding and van der Waals contacts. Consequently, the dye is firmly anchored to the collagen, so it resists being washed out during subsequent processing steps. Moreover, picric acid also suppresses non-specific binding of Sirius Red to other tissue components, ensuring that the stain is largely confined to collagen fibers.^17^ Under standard microscopy, this produces highly specific and stable red staining of collagen bundles against a pale yellow background, providing excellent contrast for visualization.

Thus, this paper aims to present a step-by-step protocol that combines Alcian Blue and Picrosirius Red stains on slides from CRC patient tumor tissue. Although combinations of different stains are common in histology, including H&E and Movat’s Pentachrome, the combination of Alcian Blue and Picrosirius Red has not been systematically applied to CRC, nor have its optical interactions been investigated. Integrating both stains on a single tissue section conserves patient tissue material, reduces imaging time, and delivers a unified spatial map of the ECM architecture of the TME. Structural importance of collagen is well documented, yet the spatial distribution of GAGs and their interplay with collagen remain under-explored, particularly in CRC where the ECM undergoes profound remodeling. This combination of stains can help address these questions and further our understanding of CRC.

## 2. MATERIALS AND METHODS

Research papers generally tend to exercise extreme brevity in this section due to a variety of reasons including word and page limitations. Nevertheless, a detailed outline of materials and methods remains essential for scientific rigor and reproducibility. This is of particular significance in cellular pathology and laboratory analyses related to solid tumor tissue. Thus, this protocol paper provides an in-depth description of materials and methods for consistency and reproducibility. Through detailed documentation of each step and corresponding materials, we offer a standardized framework that can be implemented across a broad array of laboratory environments.

### 2.1. Before you begin

#### Permissions

Patient permission for the tumor tissue study should be obtained. Tumor tissue specimens were collected from patients who underwent surgery at University of Virginia (UVA) Health, located in Charlottesville, Virginia. Biospecimens reviewed internally to ensure de-identification. All patients provided informed consent for the collection and donation of the specimens.

### 2.2. Key Resources

Key equipment and resources needed for this protocol are listed in Table 1.

**Table 1.**
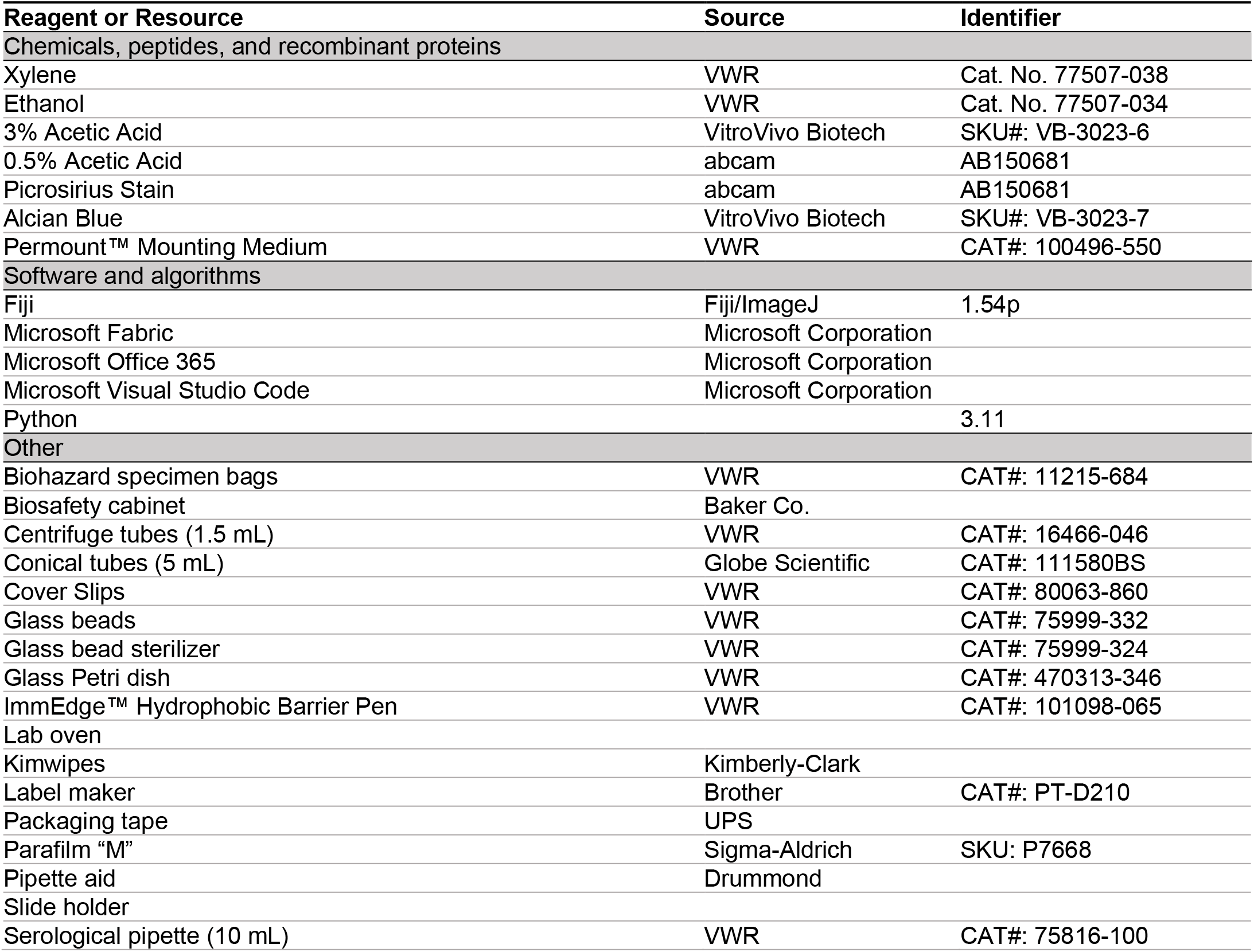
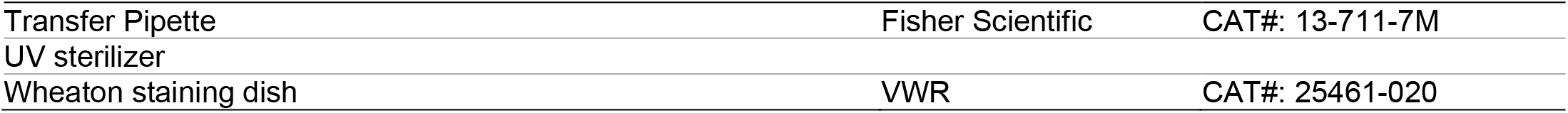
Key equipment and resources table.

### 2.3. Step-By-Step Method Details

#### 2.3.1. Alcian Blue Staining Procedure

This segment of the protocol describes steps involved in staining histology slides to visualize GAGs using Alcian Blue at pH 2.5. Strict personal and environmental safety protocols must also be followed during this protocol, including the use of appropriate personal protective equipment such as gloves, safety goggles, and respirators.

1. Label two Wheaton staining dishes as “Xylene I” and “Xylene II.”
  a. Carefully dispense 200 mL of Xylene into each dish. Promptly cover the dish with glass container.
2. Label two Wheaton staining dishes as “Ethanol 100% (a) “and “Ethanol 100% (b).”
  a. Carefully dispense 200 mL of 100% ethanol into each dish. Promptly cover the dish with glass container.
3. Label two Wheaton staining dishes as “Ethanol 95% (a)” and “Ethanol 95% (b).”
  a. Carefully dispense 200 mL of 95% ethanol into each dish. Promptly cover the dish with glass container.
4. Bake slides in laboratory oven at 65°C for 30 minutes. Increase bake time to at least 4 hours for freshly prepared slides. Let slides acclimate back to room temperature.
5. Incubate slides in Xylene.
  a. Incubate slides in Wheaton staining dish labeled “Xylene I” for 2 minutes.
  b. Immediately move slides to Wheaton staining dish labeled “Xylene II” for another 2 minutes.
6. Rehydrate slides in ethanol.
  a. Move slides to Wheaton staining dish labeled “Ethanol 100% (a)” for 2 minutes.
  b. Then to Wheaton staining dish labeled “Ethanol 100% (b)” for 2 minutes.
  c. Move slides to Wheaton staining dish labeled “Ethanol 95% (a)” for 2 minutes.
  d. Then to Wheaton staining dish labeled “Ethanol 95% (b)” for 2 minutes.
7. Rinse in distilled water for 5 minutes.
8. Draw circle around tumor section on slides with hydrophobic pen.
  a. Leave adequate space between tissue section and circumference cast by the hydrophobic pen.
  b. For the rest of this procedure, the area inside this circumference will be referred to as “Stain Area”
  c. Note: This is to ensure that stain will remain localized over the tumor tissue section on the slide. Our approach of using a hydrophobic pen coupled with smaller quantity of stain, instead of dipping entire slides in stain, is to reduce waste and environmental footprint.
9. Gently dispense adequate 3% Acetic Acid solution to cover “Stain Area” of slides. Incubate slides for 3 minutes.
10. Discard the Acetic Acid solution from the slides but do not rinse the slides.
11. Gently dispense adequate Alcian Blue solution to cover “Stain Area” of slides. Incubate for 30 minutes.
12. Rinse sections briefly in 3% Acetic Acid solution to remove Alcain blue stain from slides.
13. Wash slides in running tap water for 10 minutes. Use intermediary flow tanks to reduce water waste.
14. Rinse slides in distilled water for 5 minutes.
15. Rehydrate slides in ethanol.
  a. Move slides to Wheaton staining dish labeled “Ethanol 95% (b)” for 30 seconds.
  b. Then to Wheaton staining dish labeled “Ethanol 95% (a)” for 30 seconds.
  c. Move slides to Wheaton staining dish labeled “Ethanol 100% (b)” for 30 seconds.
  d. Then to Wheaton staining dish labeled “Ethanol 100% (a)” for 30 seconds.
16. Rinse slides in Wheaton staining dish labeled “Xylene II” for another 90 seconds (or until wax ring is gone)
17. Mount and cover slide
  a. Draw a line with Permount to mark the edge where cover slip will be placed on the slide.
  b. Gently place cover slip edge over this line of mounting media.
  c. Allow coverslip to slowly spread the mounting media over the entire tissue section and towards the other edge of the cover slip on the slide
18. Safely store Xylene.
  a. Gently transfer Xylene from Wheaton staining dish labeled “Xylene I” to an appropriate airtight container labeled “Xylene I” for use later. Log the date and number of slides to monitor duty cycles of Xylene.
  b. Gently transfer Xylene from Wheaton staining dish labeled “Xylene II” to an appropriate airtight container labeled “Xylene II” for use later. Log the date and number of slides to monitor duty cycles of Xylene.
19. Safely store Ethanol.
  a. Gently transfer 100% Ethanol from Wheaton staining dishes labeled “Ethanol 100% (a)” and “Ethanol 100% (b)” to appropriate airtight containers labeled “Ethanol 100% (a)” and “Ethanol 100% (b)” for use later. Log the date and number of slides to monitor duty cycles of 100% Ethanol.
  b. Gently transfer 95% Ethanol from Wheaton staining dishes labeled “Ethanol 95% (a)” and “Ethanol 95% (b)” to appropriate airtight containers labeled “Ethanol 95% (a)” and “Ethanol 95% (b)” for use later. Log the date and number of slides to monitor duty cycles of 95% Ethanol.

#### 2.3.2. Picrosirius Red Staining Procedure

This segment of the protocol describes steps involved in staining histology slides to visualize collagen using Picrosirius Red. Strict personal and environmental safety protocols must also be followed during this protocol, including the use of appropriate personal protective equipment such as gloves, safety goggles, and respirators.

1. Label two Wheaton staining dishes as “Xylene I” and “Xylene II.”
  a. Carefully dispense 200 mL of Xylene into each dish. Promptly cover the dish with glass container.
2. Label two Wheaton staining dishes as “Ethanol 100% (a) “and “Ethanol 100% (b).”
  a. Carefully dispense 200 mL of 100% ethanol into each dish. Promptly cover the dish with glass container.
3. Label two Wheaton staining dishes as “Ethanol 95% (a)” and “Ethanol 95% (b).”
  a. Carefully dispense 200 mL of 95% ethanol into each dish. Promptly cover the dish with glass container.
4. Bake slides in laboratory oven at 65°C for 30 minutes. Increase bake time to at least 4 hours for freshly prepared slides. Let slides acclimate back to room temperature.
5. Incubate slides in Xylene.
  a. Incubate slides in Wheaton staining dish labeled “Xylene I” for 2 minutes.
  b. Immediately move slides to Wheaton staining dish labeled “Xylene II” for another 2 minutes.
6. Rehydrate slides in ethanol.
  a. Move slides to Wheaton staining dish labeled “Ethanol 100% (a)” for 2 minutes.
  b. Then to Wheaton staining dish labeled “Ethanol 100% (b)” for 2 minutes.
  c. Move slides to Wheaton staining dish labeled “Ethanol 95% (a)” for 2 minutes.
  d. Then to Wheaton staining dish labeled “Ethanol 95% (b)” for 2 minutes.
7. Rinse in distilled water for 5 minutes.
8. Draw circle around tumor section on slides with hydrophobic pen.
  a. Leave adequate space between tissue section and circumference cast by the hydrophobic pen.
  b. For the rest of this procedure, the area inside this circumference will be referred to as “Stain Area”
  c. Note: This is to ensure that stain will remain localized over the tumor tissue section on the slide. Our approach of using a hydrophobic pen coupled with smaller quantity of stain, instead of dipping entire slides in stain, is to reduce waste and environmental footprint.
9. Gently dispense adequate Picrosirius Red Solution to cover “Stain Area” of slides with transfer pipette. Incubate for 60 minutes.
10. Rinse slides quickly in 2 changes of 0.5% Acetic Acid Solution
11. Rehydrate slides in ethanol.
  a. Move slides to Wheaton staining dish labeled “Ethanol 95% (b)” for 30 seconds.
  b. Then to Wheaton staining dish labeled “Ethanol 95% (a)” for 30 seconds.
  c. Move slides to Wheaton staining dish labeled “Ethanol 100% (b)” for 30 seconds.
  d. Then to Wheaton staining dish labeled “Ethanol 100% (a)” for 30 seconds.
12. Rinse slides in Wheaton staining dish labeled “Xylene II” for another 90 seconds (or until wax ring is gone)
13. Mount and cover slide
  a. Draw a line with Permount to mark the edge where cover slip will be placed on the slide.
  b. Gently place cover slip edge over this line of mounting media.
  c. Allow coverslip to slowly spread the mounting media over the entire tissue section and towards the other edge of the cover slip on the slide
14. Safely store Xylene.
  a. Gently transfer Xylene from Wheaton staining dish labeled “Xylene I” to an appropriate airtight container labeled “Xylene I” for use later. Log the date and number of slides to monitor duty cycles of Xylene.
  b. Gently transfer Xylene from Wheaton staining dish labeled “Xylene II” to an appropriate airtight container labeled “Xylene II” for use later. Log the date and number of slides to monitor duty cycles of Xylene.
15. Safely store Ethanol.
  c. Gently transfer 100% Ethanol from Wheaton staining dishes labeled “Ethanol 100% (a)” and “Ethanol 100% (b)” to appropriate airtight containers labeled “Ethanol 100% (a)” and “Ethanol 100% (b)” for use later. Log the date and number of slides to monitor duty cycles of 100% Ethanol.
  d. Gently transfer 95% Ethanol from Wheaton staining dishes labeled “Ethanol 95% (a)” and “Ethanol 95% (b)” to appropriate airtight containers labeled “Ethanol 95% (a)” and “Ethanol 95% (b)” for use later. Log the date and number of slides to monitor duty cycles of 95% Ethanol.

#### 2.3.3. Dual-Stain – Alcian Blue and Picrosirius Red – Staining Procedure

This segment of the protocol describes steps involved in dual-staining histology slides to simultaneously visualize GAGs and collagen using Alcian Blue and Picrosirius Red. Strict personal and environmental safety protocols must also be followed during this protocol, including the use of appropriate personal protective equipment such as gloves, safety goggles, and respirators.

1. Label two Wheaton staining dishes as “Xylene I” and “Xylene II.”
  a. Carefully dispense 200 mL of Xylene into each dish. Promptly cover the dish with glass container.
2. Label two Wheaton staining dishes as “Ethanol 100% (a) “and “Ethanol 100% (b).”
  a. Carefully dispense 200 mL of 100% ethanol into each dish. Promptly cover the dish with glass container.
3. Label two Wheaton staining dishes as “Ethanol 95% (a)” and “Ethanol 95% (b).”
  a. Carefully dispense 200 mL of 95% ethanol into each dish. Promptly cover the dish with glass container.
4. Bake slides in laboratory oven at 65°C for 30 minutes. Increase bake time to at least 4 hours for freshly prepared slides. Let slides acclimate back to room temperature.
5. Incubate slides in Xylene.
  a. Incubate slides in Wheaton staining dish labeled “Xylene I” for 2 minutes.
  b. Immediately move slides to Wheaton staining dish labeled “Xylene II” for another 2 minutes.
6. Rehydrate slides in ethanol.
  a. Move slides to Wheaton staining dish labeled “Ethanol 100% (a)” for 2 minutes.
  b. Then to Wheaton staining dish labeled “Ethanol 100% (b)” for 2 minutes.
  c. Move slides to Wheaton staining dish labeled “Ethanol 95% (a)” for 2 minutes.
  d. Then to Wheaton staining dish labeled “Ethanol 95% (b)” for 2 minutes.
7. Rinse in distilled water for 5 minutes.
8. Draw circle around tumor section on slides with hydrophobic pen.
  a. Leave adequate space between tissue section and circumference cast by the hydrophobic pen.
  b. For the rest of this procedure, the area inside this circumference will be referred to as “Stain Area”
  c. Note: This is to ensure that stain will remain localized over the tumor tissue section on the slide. Our approach of using a hydrophobic pen coupled with smaller quantity of stain, instead of dipping entire slides in stain, is to reduce waste and environmental footprint.
9. Gently dispense adequate 3% Acetic Acid solution to cover “Stain Area” of slides with transfer pipette. Incubate slides for 3 minutes.
10. Discard the Acetic Acid solution from the slides but do not rinse the slides.
11. Gently dispense adequate Alcian Blue solution to cover “Stain Area” of slides. Incubate for 20 minutes.
12. Rinse sections briefly in 3% Acetic Acid solution to remove Alcain blue stain from slides.
13. Wash slides in running tap water for 10 minutes. Use intermediary flow tanks to reduce water waste.
14. Rinse slides in distilled water for 5 minutes.
15. Gently dispense adequate Picrosirius Red Solution to cover “Stain Area” of slides. Incubate for 60 minutes.
16. Rinse slides quickly in 2 changes of 0.5% Acetic Acid Solution
17. Rehydrate slides in ethanol.
  a. Move slides to Wheaton staining dish labeled “Ethanol 95% (b)” for 30 seconds.
  b. Then to Wheaton staining dish labeled “Ethanol 95% (a)” for 30 seconds.
  c. Move slides to Wheaton staining dish labeled “Ethanol 100% (b)” for 30 seconds.
  d. Then to Wheaton staining dish labeled “Ethanol 100% (a)” for 30 seconds.
18. Rinse slides in Wheaton staining dish labeled “Xylene II” for another 90 seconds (or until wax ring is gone)
19. Mount and cover slide
  a. Draw a line with Permount to mark the edge where cover slip will be placed on the slide.
  b. Gently place cover slip edge over this line of mounting media.
  c. Allow coverslip to slowly spread the mounting media over the entire tissue section and towards the other edge of the cover slip on the slide
20. Safely store Xylene.
  a. Gently transfer Xylene from Wheaton staining dish labeled “Xylene I” to an appropriate airtight container labeled “Xylene I” for use later. Log the date and number of slides to monitor duty cycles of Xylene.
  b. Gently transfer Xylene from Wheaton staining dish labeled “Xylene II” to an appropriate airtight container labeled “Xylene II” for use later. Log the date and number of slides to monitor duty cycles of Xylene.
21. Safely store Ethanol.
  a. Gently transfer 100% Ethanol from Wheaton staining dishes labeled “Ethanol 100% (a)” and “Ethanol 100% (b)” to appropriate airtight containers labeled “Ethanol 100% (a)” and “Ethanol 100% (b)” for use later. Log the date and number of slides to monitor duty cycles of 100% Ethanol.
  b. Gently transfer 95% Ethanol from Wheaton staining dishes labeled “Ethanol 95% (a)” and “Ethanol 95% (b)” to appropriate airtight containers labeled “Ethanol 95% (a)” and “Ethanol 95% (b)” for use later. Log the date and number of slides to monitor duty cycles of 95% Ethanol.

#### 2.3.4. Image Processing and Estimating Collagen-GAG ECM Contribution

This segment of the protocol describes steps involved in digital processing of dual-stained histology slides to quantify collagen and GAG distribution using Fiji^19^ software. This workflow generates quantitative spatial maps showing the distribution relationship between collagen and GAG components in the tumor microenvironment. Ensure adequate computer processing power and storage space for handling large TIFF image files.

1. Acquire high-resolution images using Lionheart microscope at 20X magnification.
  a. Set image format to lossless RGB TIFF to preserve all spectral information.
  b. Ensure consistent illumination and white balance across all images.
  c. Save images with systematic naming convention including sample ID and magnification.
2. Open Fiji software
  a. Install Color Deconvolution 2 plugin if not already available.
  b. Navigate to Plugins > Color Deconvolution 2 to verify installation.
  c. If not installed, download from ImageJ plugin repository and restart Fiji.
3. Load dual-stained image into Fiji.
  a. Open RGB TIFF file using File > Open.
  b. Verify image displays correctly with both red (collagen) and blue (GAG) staining visible.
4. Initiate color deconvolution process.
  a. Navigate to Plugins > Color Deconvolution 2.
  b. Select “From ROI” from dropdown menu for Vectors.
  c. Select “8bit_Transmittance” from dropdown menu for Output.
5. Select Region of Interest (ROI) for red stain calibration.
  a. Use Rectangle tool to select 50×50 to 100×100 pixel area with dense, uniform Picrosirius Red staining.
  b. Avoid tissue folds, artifacts, or areas with mixed staining.
  c. Choose regions with intense, pure red collagen staining.
  d. Click “Pick [1st Stain]” button and select the ROI.
6. Select Region of Interest (ROI) for blue stain calibration.
  a. Use Rectangle tool to select similar-sized area with strong, uniform Alcian Blue staining.
  b. Avoid overlapping regions where both stains might be present.
  c. Ensure pure blue GAG staining without background interference.
  d. Click “Pick [2nd Stain]” button and select the ROI.
7. Select Region of Interest (ROI) for background calibration.
  a. Use Rectangle tool to select unstained tissue area or slide background.
  b. Avoid debris, dust, or optical artifacts.
  c. Choose area representing true negative staining.
  d. Click “Pick [3rd Stain]” button and select the ROI.
8. Execute color deconvolution.
  a. Click “OK” to run deconvolution algorithm.
  b. Three separate grayscale images will be generated: Colour_1 (red component), Colour_2 (blue component), and Colour_3 (background).
9. Convert deconvolved images to 8-bit format.
  a. Select Colour_1 image window.
  b. Navigate to Image > Type > 8-bit.
  c. Repeat for Colour_2 and Colour_3 images.
10. Apply Otsu auto-threshold to create binary masks.
  a. Select Colour_1 image (red/collagen component).
  b. Navigate to Image > Adjust > Threshold.
  c. Select “Otsu” from algorithm dropdown menu.
  d. Click “Apply” to create binary mask where white pixels represent stained areas.
  e. Repeat steps a-d for Colour_2 image (blue/GAG component).
11. Measure percentage area stained for each component.
  a. Navigate to Analyze > Set Measurements and ensure “Area fraction” is checked.
  b. Select binary mask for Colour_1 (collagen).
  c. Use Analyze > Measure to obtain percentage area stained.
  d. Record result as “% Collagen Area.”
  e. Repeat steps b-d for Colour_2 binary mask and record as “% GAG Area.”
12. Save processed images and measurements.
  a. Save binary masks as TIFF files with descriptive names (e.g., “SampleID_Collagen_Binary.tif”).
  b. Export measurement results to Excel or CSV format using File > Save As.
  c. Create systematic file organization structure for analysis tracking.
13. Document processing parameters for reproducibility.
  a. Record ROI coordinates and sizes used for color deconvolution.
  b. Note any manual threshold adjustments if Otsu auto-threshold was modified.
  c. Log image acquisition settings and processing software versions.

## 3. RESULTS AND DISCUSSION

To assess fidelity of our dual-staining approach, we compared ECM component detection between single-stain Alcian Blue and single-stain Picrosirius Red versus the combined dual-stain protocol on serial sections from the same CRC patient tumor (**Fig. 1**). Visual examination shows that the dual-stain protocol successfully preserved the distinct blue coloration of GAGs and red coloration of collagen without apparent spectral interference or loss of staining intensity. Furthermore, the dual staining protocol with distinct coloration of each component enabled visualization and characterization of ECM spatial organization within the TME. The slides were then scanned for quantification and analysis. All images were obtained using a 20X objective in color brightfield.

**Figure 1.**
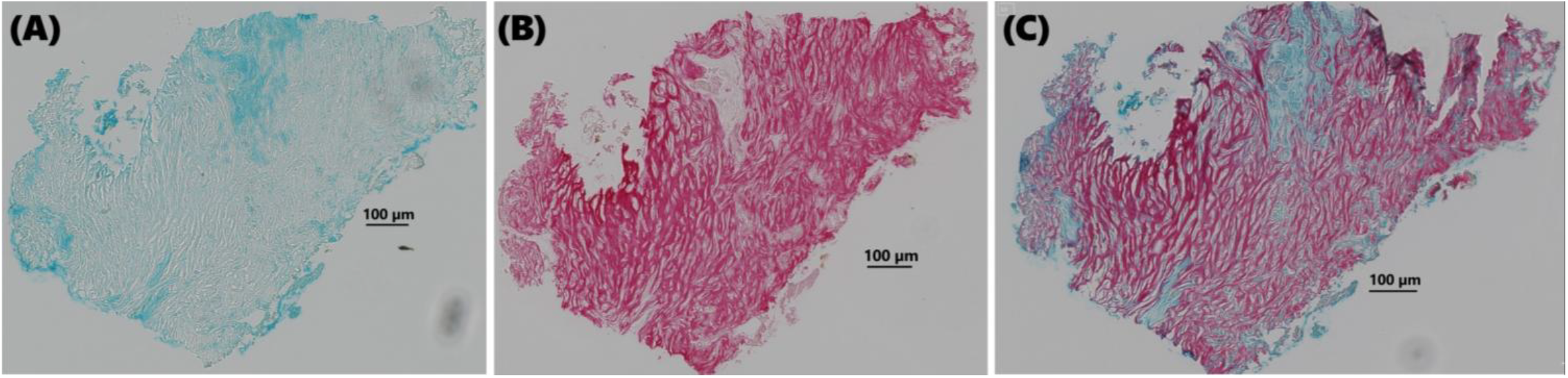
Comparison of single and dual-staining approaches in CRC tissue. Representative images from the same patient tumor showing **(A)** Alcian Blue single-stain highlighting GAGs in blue, **(B)** Picrosirius Red single-stain highlighting collagen in red, and **(C)** dual-stain combining both Alcian Blue and Picrosirius Red on the same tissue section. All images acquired in color brightfield at 20X magnification. Scale bars = 100 μm.

Digital color deconvolution analysis using the *Color Deconvolution 2* plugin in Fiji^19^ provided quantitative assessment of the dual-staining approach (**Fig. 2**). When comparing the blue component deconvolved from the Alcian Blue single-stain (**Fig. 2A**) with the blue component extracted from the dual-stain image (**Fig. 2B**), we observed remarkably similar GAG distribution patterns despite the different staining protocols and tissue sections albeit from the same patient tumor. The spatial organization of GAG-rich regions was faithfully preserved in the dual-stain approach, with comparable staining intensity and morphological detail. Similarly, comparison of the red component from Picrosirius Red single-stain (**Fig. 2C**) with the red component from dual-stain deconvolution (**Fig. 2D**) demonstrated excellent preservation of collagen fiber architecture and distribution.

**Figure 2.**
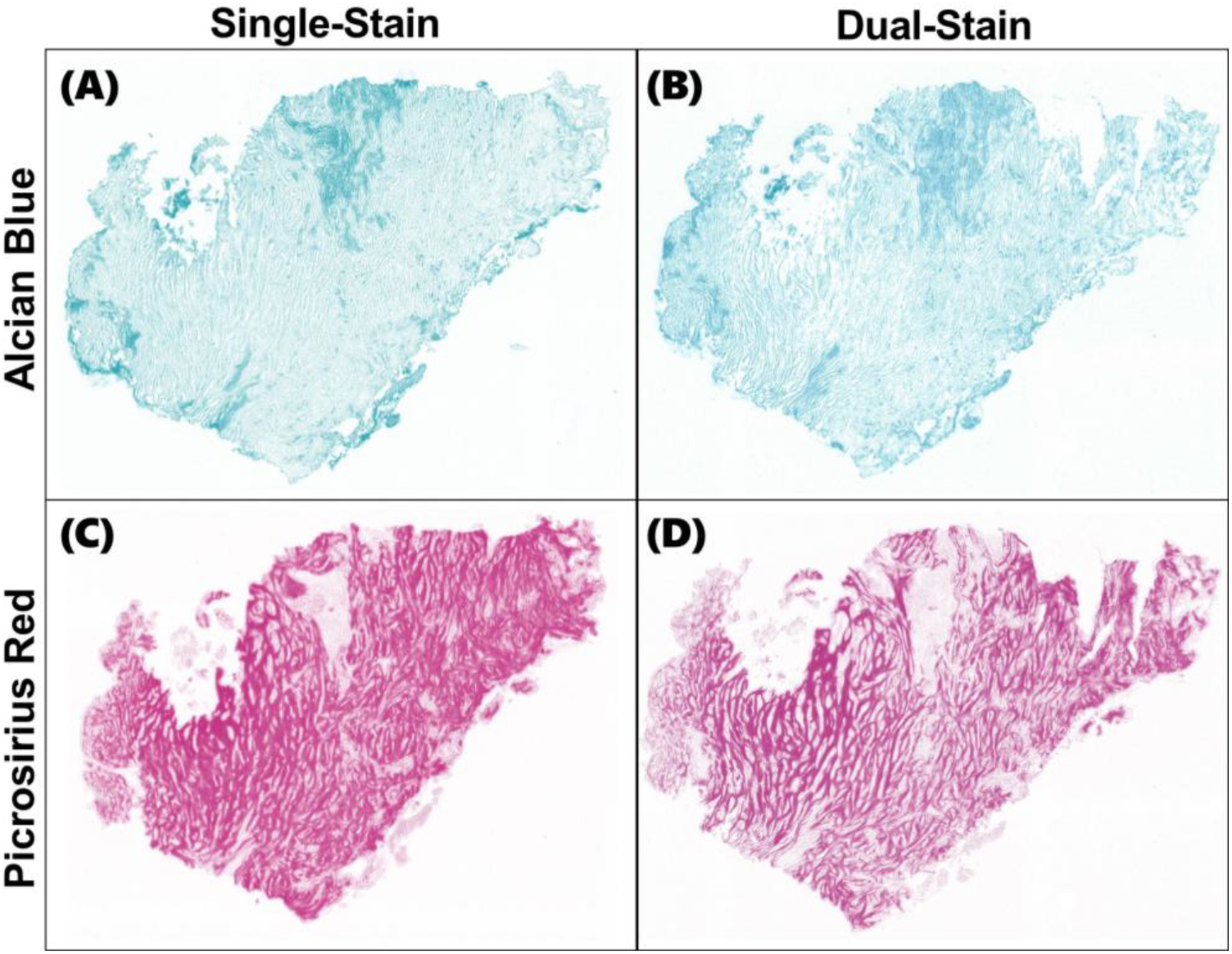
Digital color deconvolution. Deconvolved color components comparing single-stain and dual-stain approaches. **(A)** Blue component deconvolved from Alcian Blue single-stain image from Fig. 1A. **(B)** Blue component deconvolved from dual-stain image from Fig. 1C, demonstrating closely preserved GAG detection. **(C)** Red component deconvolved from Picrosirius Red single-stain image from Fig. 1B. **(D)** Red component deconvolved from dual-stain image from Fig. 1C, demonstrating closely preserved collagen detection. Despite different tissue sections and staining protocols, dual-stain deconvolution closely approximates single-stain results for both ECM components.

Binary threshold analysis further confirmed the quantitative fidelity of the dual-staining protocol (**Fig. 3**). Otsu auto-threshold application to deconvolved images generated binary masks that accurately captured the spatial extent of ECM component staining. The GAG masks derived from single-stain (**Fig. 3A**) and dual-stain (**Fig. 3B**) approaches showed consistent identification of GAG-rich regions, while collagen masks from single-stain (**Fig. 3C**) and dual-stain (**Fig. 3D**) protocols demonstrated similar detection of fibrillar collagen networks.

**Figure 3.**
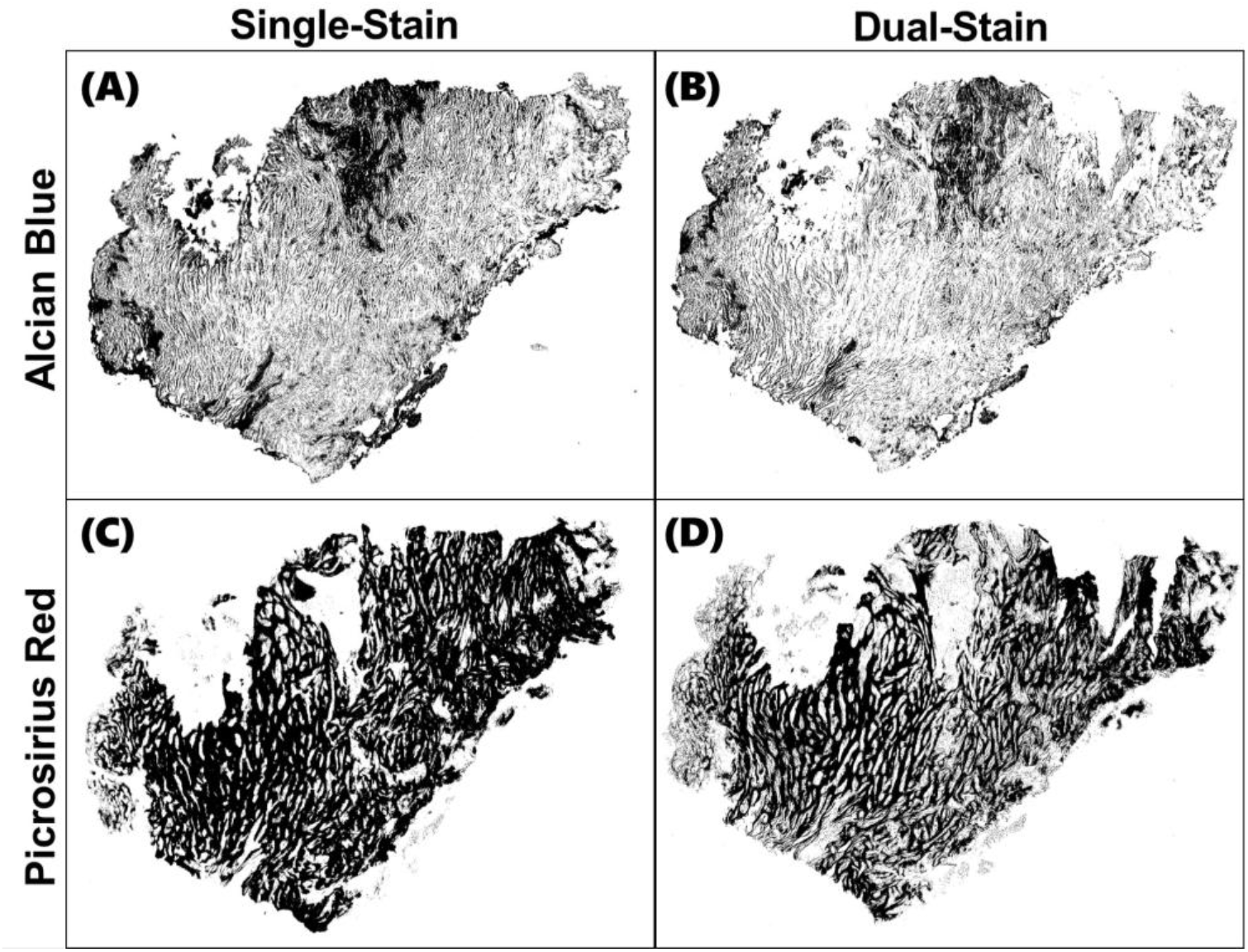
Binary threshold analysis. Otsu auto-threshold applied to deconvolved images to create binary masks for quantification. **(A)** Thresholded GAG mask from Alcian Blue single-stain. **(B)** Thresholded GAG mask from dual-stain deconvolution. **(C)** Thresholded collagen mask from Picrosirius Red single-stain. **(D)** Thresholded collagen mask from dual-stain deconvolution. Binary masks demonstrate that dual-staining preserves the spatial distribution patterns and relative abundance of both ECM components compared to individual staining protocols.

These results demonstrate that our dual-staining protocol maintains the specificity and sensitivity of individual staining methods while enabling simultaneous ECM component characterization and spatial organization. The preserved spatial fidelity and quantitative accuracy support the use of this approach for comprehensive TME analysis, offering significant advantages in terms of tissue conservation, processing efficiency, and direct spatial comparison of collagen and GAG distribution patterns within the same histological section. The protocol’s primary advantage lies in enhancing spatial and quantitative analysis of the TME, particularly when tissue sections with intact ECM are limited. This is especially valuable for tissue sites that are not easily accessible, or for downstream research applications with no direct benefit to the patient to justify obtaining additional tissue. Furthermore, the spectral clarity of the distinct blue and red wavelengths from our dual-stain protocol reduces the ambiguity inherent in multi-component stains with overlapping color ranges. Unlike traditional Movat’s pentachrome methods that produce yellow-to-red gradients encompassing multiple tissue components, our dual-stain approach provides unambiguous component identification essential for both visual assessment and digital analysis workflows.

## 4. CONCLUSION

We have developed and validated a robust dual-staining protocol that simultaneously visualizes and quantifies collagen and GAG distribution in the CRC TME. This approach addresses a gap in ECM characterization methodology by combining the specificity of traditional single-stain techniques with the efficiency and spatial fidelity required for comprehensive microenvironment analysis. The protocol’s validation demonstrates concordance between dual-stain and single staining approaches, confirming that simultaneous ECM component detection does not significantly compromise accuracy or sensitivity of either Alcian Blue or Picrosirius Red staining. The clear spectral separation achieved through our blue-red chromogen combination eliminates the ambiguity inherent in overlapping multi-component stains, making this approach particularly well-suited for digital pathology workflows and automated image analysis systems.

Beyond methodological advancement, this protocol directly addresses pressing translational research needs. As tissue availability is limited for downstream research applications, our approach maximizes the analytical value extracted from precious patient specimens. The standardized nature of the protocol, utilizing common histology equipment and reagents, ensures broad accessibility and reproducibility across diverse laboratory settings.

The timing of this methodology is also significant given the regulatory shift toward human-relevant preclinical models and NAMs. Our protocol adds to the tools needed to characterize and validate the ECM fidelity of patient-derived models ensuring that these next-generation platforms can better approximate ECM conditions that drive disease progression and therapeutic response. This capability is essential for bridging the translational gap that continues to hinder cancer drug development and contributes to high failure rates in clinical trials.

The adaptability of this approach extends beyond CRC. The fundamental principles of simultaneous collagen-GAG visualization are applicable across solid tumor types, opening new avenues for comparative ECM analysis in pancreatic, ovarian, breast, lung, and other cancers in which stromal remodeling drives progression and treatment resistance. Furthermore, the protocol’s compatibility with complementary techniques, including advanced imaging modalities, positions it as a useful method for integrated TME characterization.

As precision medicine increasingly recognizes the TME as a therapeutic target and prognostic indicator, standardized approaches for ECM characterization will become indispensable. This protocol provides the cancer researchers with a validated, step-by-step, accessible tool for ECM analysis. We hope that this methodology will accelerate discoveries linking stromal architecture to cancer biology and contribute to improved patient outcomes.

## DECLARATIONS

### Author Contributions

Conceptualization, KIB; methodology, LAB, SJ, DS, KKB, BKB, and KIB; validation, CLC, and AT; formal analysis, LAB, SJ, and DS; investigation, LAB, SJ, DS, KKB, and BKB; resources, AT and KIB; data curation, LAB, SJ, DS, KKB, and BKB; writing—original draft preparation, LAB, SJ, DS, KKB, BKB, and KIB; writing—review and editing, All authors; supervision, AT and KIB; project administration, CLC, KIB; funding acquisition, KIB. All authors have read and agreed to the published version of the manuscript.

### Funding

This work was funded by CerFlux, Innovate Alabama, the National Science Foundation, grant number TI-2321805 and the National Cancer Institute at the National Institutes of Health, grant number 1R43CA254493-01.

### Data Availability Statement

Additional data available upon request.

## Acknowledgments

We thank our colleagues at the Advanced Medical Imaging Research Division at the University of Alabama at Birmingham (UAB), the UAB Pathology Core Research Lab, the Hugh Kaul Precision Medicine Institute at UAB, the O’Neal Comprehensive Cancer Center, the Alabama School of Fine Arts, and the Aga Khan University Nairobi Cancer Centre for their support and collaboration.

## Conflicts of Interest

Dr. Budhwani is co-inventor of issued (and pending) patents pertaining to *in vitro, ex vivo*, and cancer supermodel technologies.

## REFERENCES

1. Morabito M, Thibodot P, Gigandet A, et al. Liver Extracellular Matrix in Colorectal Liver Metastasis. Cancers (Basel) 2025;17(6):1–28.

2. American Cancer Society. Colorectal Cancer Facts & Figures 2023-2025. 2023.

3. Siegel RL, Giaquinto AN, Jemal A. Cancer statistics, 2024. CA Cancer J Clin [Internet] 2024;74(1):12–49. Available from: https://acsjournals.onlinelibrary.wiley.com/doi/10.3322/caac.21820

4. Lee JJ, Ng KY, Bakhtiar A. Extracellular matrix: unlocking new avenues in cancer treatment. Biomark Res 2025;13(1):1–25.

5. Liang Y, Lv Z, Huang G, et al. Prognostic significance of abnormal matrix collagen remodeling in colorectal cancer based on histologic and bioinformatics analysis. Oncol Rep 2020;44(4):1671–85.

6. Morla S. Glycosaminoglycans and glycosaminoglycan mimetics in cancer and inflammation. Int J Mol Sci 2019;20(8).

7. Huang J, Zhang L, Wan D, et al. Extracellular matrix and its therapeutic potential for cancer treatment. Signal Transduct Target Ther [Internet] 2021;6(1). Available from: 10.1038/s41392-021-00544-0

8. Thomasy H. Is This the End of Animal Testing ? FDA Announces Plans to Phase Out Animals in Drug Safety Studies. Sci [Internet] 2025;1–10. Available from: https://www.the-scientist.com/is-this-the-end-of-animal-testing-fda-announces-plans-to-phase-out-animals-in-drug-safety-studies-73031

9. Budhwani KI, Patel ZH, Guenter RE, Charania AA. A hitchhiker’s guide to cancer models. Trends Biotechnol [Internet] 2022;1–13. Available from: 10.1016/j.tibtech.2022.04.003

10. U.S. Food and Drug Administration. FDA Announces Plan to Phase Out Animal Testing Requirement for Monoclonal Antibodies and Other Drugs [Internet]. 2025. Available from: https://www.fda.gov/news-events/press-announcements/fda-announces-plan-phase-out-animal-testing-requirement-monoclonal-antibodies-and-other-drugs

11. National Institutes of Health. Catalyzing The Development and Use of Novel Alternative Methods. 2024;(December).

12. Budhwani KI, Bortone K, Menon S, Scully M, Mortko CJ. From Discovery to Deals: Bridging the Valley-of-Death with a Molecule-to-Medicine Expressway [Internet]. In: BIO International Convention. 2025. p. 1–4.Available from: https://convention.bio.org/program-1/discovery-deals-bridging-valley-death-molecule-medicine-expressway

13. Mesa N. FDA, NIH Accelerate Shift Away From Animal Research as Experts Warn of Limitations. BioSpace [Internet] 2025;1–5. Available from: https://www.biospace.com/drug-development/fda-nih-accelerate-shift-away-from-animal-research-as-experts-warn-of-limitations

14. VulcanChem. Alcian blue Chemical Structure and Properties. 2025;5–9.

15. Gandhi NS, Mancera RL. The structure of glycosaminoglycans and their interactions with proteins. Chem Biol Drug Des 2008;72(6):455–82.

16. BenchChem. Sirius red F3B hexaanion [Internet]. 2025;1–12. Available from: https://www.benchchem.com/es/product/b1239346

17. Arun Gopinathan P, Kokila G, Siddeeqh S, Prakash R, L P. Reexploring picrosirius red: A review. Indian J Pathol Oncol 2020;7(2):196–203.

18. Lattouf R, Younes R, Lutomski D, et al. Picrosirius Red Staining: A Useful Tool to Appraise Collagen Networks in Normal and Pathological Tissues. J Histochem Cytochem 2014;62(10):751–8.

19. Schindelin J, Arganda-Carreras I, Frise E, et al. Fiji: An open-source platform for biological-image analysis. Nat Methods 2012;9(7):676–82.

